# Rapid Tilt-Series Acquisition for Electron Cryotomography

**DOI:** 10.1101/454587

**Authors:** Georges Chreifi, Songye Chen, Lauren Ann Metskas, Mohammed Kaplan, Grant J. Jensen

## Abstract

Using a new Titan Krios stage equipped with a single-axis holder, we developed two methods to accelerate the collection of tilt-series. We demonstrate a continuous-tilting method that can record a tilt-series in seconds (about 100x faster than current methods), but with loss of details finer than ∼4 nm. We also demonstrate a fast-incremental method that can record a tilt-series about 10x faster than current methods and with similar resolution. We characterize the utility of both methods in real biological electron cryotomography workflows. We identify opportunities for further improvements in hardware and software and speculate on the impact such advances could have on structural biology.

## Introduction

Electron cryotomography (ECT) allows the 3D visualization of intact cells and other biological samples *in situ* using a transmission electron microscope (TEM). In traditional ECT, a frozen-hydrated biological sample is placed on a goniometric stage inside the TEM and progressively tilted to different angles relative to the imaging electron beam, typically in 1°, 2°, or 3° increments, throughout a tilt range of approximately −60° to +60°. Two-dimensional projection images are recorded by a camera at each tilt angle, producing a tilt-series. These images are then aligned and computationally merged into a 3D reconstruction of the sample, or tomogram. Because high-energy imaging electrons damage biological materials, the total electron dose is limited and tomograms have poor signal-to-noise ratio (SNR). Where more than one copy of a structure is present, their reconstructions can be averaged (subtomogram averaging) to improve SNR and reveal high-resolution detail (Murphy et al., 2006, Briggs, 2013;). ECT workflows that involve subtomogram averaging often require users to collect hundreds or even thousands of tomograms over days or weeks on the cryo-EM, making them very expensive (Chang et al., 2016; Hu et al., 2015). One major bottleneck in traditional ECT is the time taken to collect a tilt-series, typically ranging from 20-60 min. This inefficiency is due in large part to the mechanical movements of the TEM goniometer and cryoholder, which are generally not stable beyond the micrometer range. Many tomography software packages, including *SerialEM* (Mastronarde, 2005), *TOM* (Nickell et al., 2005), Tomography (Thermo Fisher), USCF Tomography (Zheng et al., 2010), Leginon (Suloway et al., 2009)), and *EM-Tomo* (TVIPS GmbH) compensate for these movements by including automated tracking and focusing steps for each tilt during data collection that electronically correct for stage shifts in x, y, and z. While these solutions help keep the target centered in the field of view, they make the process slow.

Using a new, more eucentric single-axis holder (Figure 1), here we developed two methods that accelerate tilt-series acquisition by eliminating tracking. The continuous-tilting method records movie frames while the stage is continuously tilted, resulting in a tilt-series containing thousands of extremely low SNR images. The fast-incremental method records movie frames while the stage is tilted incrementally, stopping briefly at discrete tilt angles to unblank the beam and record an image. We show that the fast-incremental method can achieve similar results as previous methods, but in a fraction of the time.

**Figure 1.**
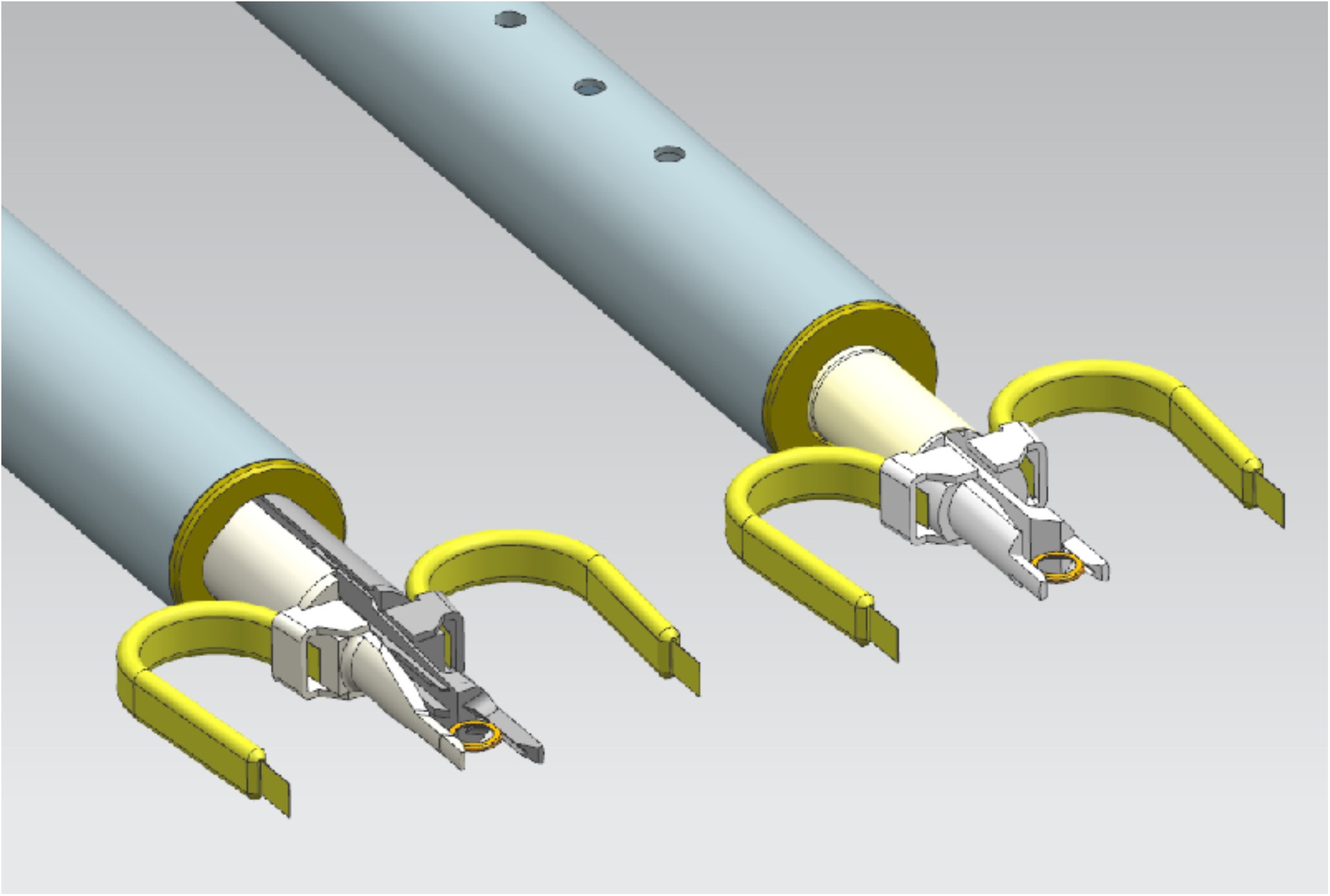
Diagram of traditional stage equipped with a dual-axis holder (left) and a stage equipped with a single-axis holder (right). Figure courtesy of Twan van den Oetelaar, Thermo Fisher Scientific.

## Results & Discussion

### Continuous-tilting method - speed

In our continuous-tilting method, the camera is directed to record a long movie while the target is exposed to the electron beam and the stage is tilted continuously. Individual images were saved at a camera framerate of 20-40 fps and at various stage tilting speeds, resulting in tilt-series containing hundreds or thousands of images (Table 1). We first assessed the speed of the continuous-tilting method by collecting tilt-series of *Bdellovibrio bacteriovorus* at several conventional pixel sizes, ranging from 4.32 Å/pixel to 1.09 Å/pixel (Table 1). In this method, the stage tilting speed is limited by the K2 camera’s optimal counting rate of 10 e^−^/pix/s. For a given target total dose (in e^−^/Å^2^) to be delivered during the tilt-series, the higher the magnification, the greater the number of camera pixels are involved, allowing a faster tilting speed. In our tilt-series of *B. bacteriovorus*, for instance, a total dose of 100 e^−^/Å^2^ was distributed in only 12 s of continuous-tilting and exposure at 1.09 Å/pixel, but 126 s were required at 4.32 Å/pixel (Table 1). Unfortunately, after the tilting and recording ended, the *SerialEM* interface remained disabled while the large movie file was transferred from the camera to the computer, preventing us from issuing any additional commands to the microscope. Therefore, instead of completing in seconds, tilt-series collected at 1.09 Å/pixel and containing 480 total images took 5.5 min to complete, and tilt-series collected at 4.32 Å/pixel and containing 1250 images took 9.7 min to complete (Table 1). As discussed below, next-generation cameras should eliminate this delay.

**Table 1.** Time taken to record tilt series using the continuous tilting method with different pixel sizes. Files were saved without gain normalization as TIFF files with LZW compression.

| Nominal Magnification | Pixel Size (Å) | Exposure time (s) | Total frames | Total time per ts (min) |
| --- | --- | --- | --- | --- |
| 33kx | 4.32 | 126 | 5040 or less | 9.7 |
| 53kx | 2.74 | 50 | 2000 or less | 7.6 |
| 81kx | 1.78 | 20 | 800 | 6.7 |
| 130kx | 1.09 | 12 | 480 | 5.5 |

### Continuous-tilting method - processing thousands of very-low-dose images

Because one of the resolution limitations in a continuous-tilting method is the arc of tilt-angles superimposed in each movie frame, we recorded images at the highest frame-rate possible with our K2 camera (40 frames/second). Further, because the sensitivity of biological samples to radiation prevented us from increasing the total dose used, each image had a very low SNR, making fiducial tracking and tilt-series alignment impossible using a conventional ECT workflow. To overcome this obstacle, we wrote a script called *Neighbor-enhance* (Figure 2) that stretches, aligns, and averages blocks of neighboring frames in a tilt-series to enhance the contrast of the fiducial markers. This allowed the gold fiducials to be automatically detected and tracked. For more details on *Neighbor-enhance*, see the Methods section.

**Figure 2.**
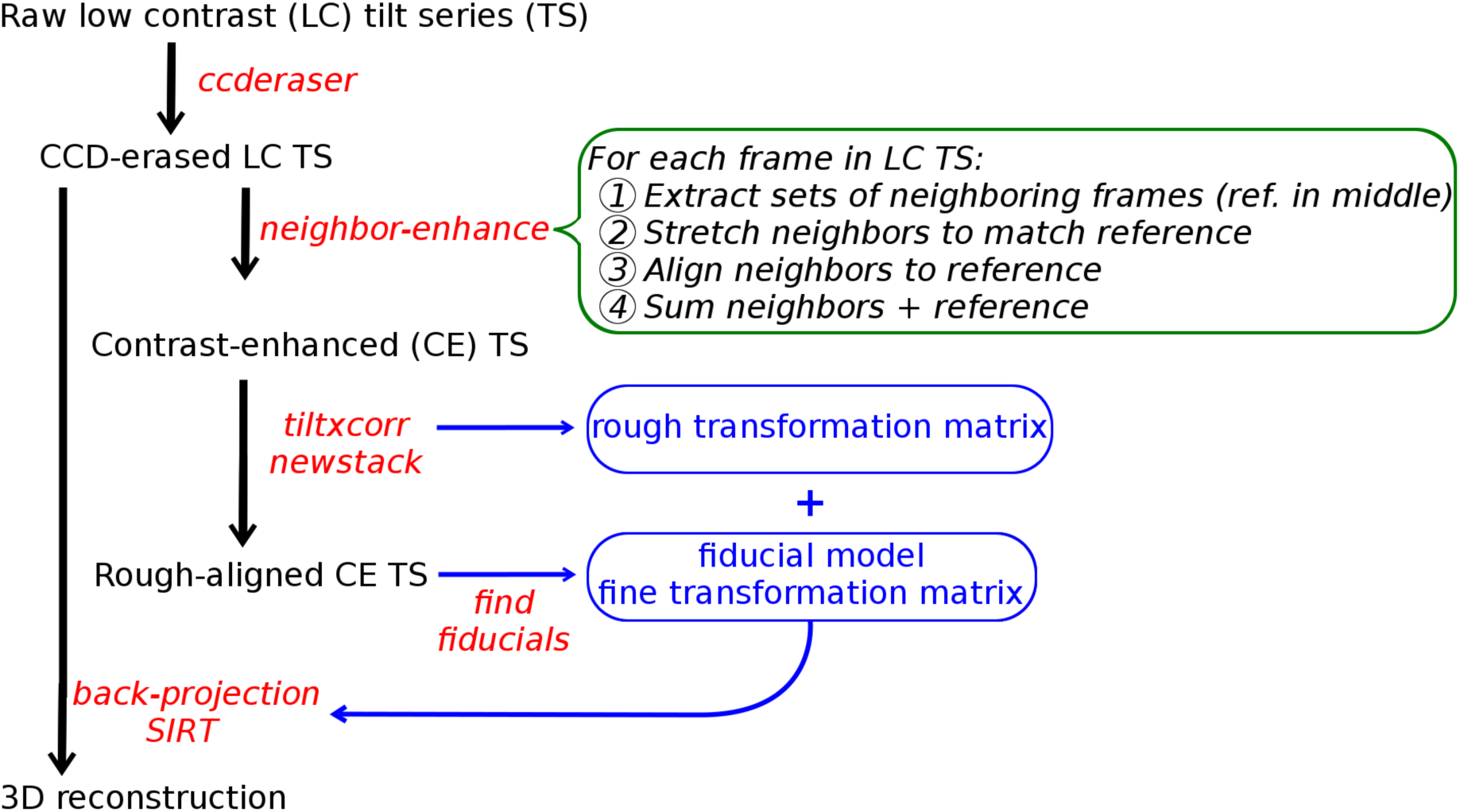
Workflow for processing data acquired using the continuous tilting method. Tilt-series data are depicted in black. Programs and scripts are depicted in red. Alignment transforms are depicted in blue. The green box contains a description of “neighbor-enhance”, an IMOD script written to enhance the contrast of the fiducial markers (see methods section for details).

### Continuous-tilting method - stage eucentricity

We characterized the eucentricity of the stage during continuous-tilting by determining the translations in X and Y required to align the projection images. We plotted these shifts in X and Y at different stage tilting speeds, from 1 °/s to 10 °/s, with the X-axis parallel to the tilt axis (Figure 3). The overall stage behavior during continuous-tilting followed a similar trend at all tilting speeds, with the majority of the movement (about 250 nm total) occurring perpendicular to the tilt axis, and very little movement along the tilt axis (about 30-75 nm total). The stage movements were more consistent at higher tilt speeds (Figure 3C and 3D). Given a shift of 150 nm in any direction, the corresponding losses of field of view on our 4k x 4k camera were less than 10% at 4.32 Å/pixel (Figure 3A), and up to about 35% at 1.09 Å/pixel (Figure 3D), suggesting that the stage is eucentric enough that tracking can be omitted at most common pixel sizes. Because even small errors in stage height increase movements, our data actually underestimate stage performance.

**Figure 3.**
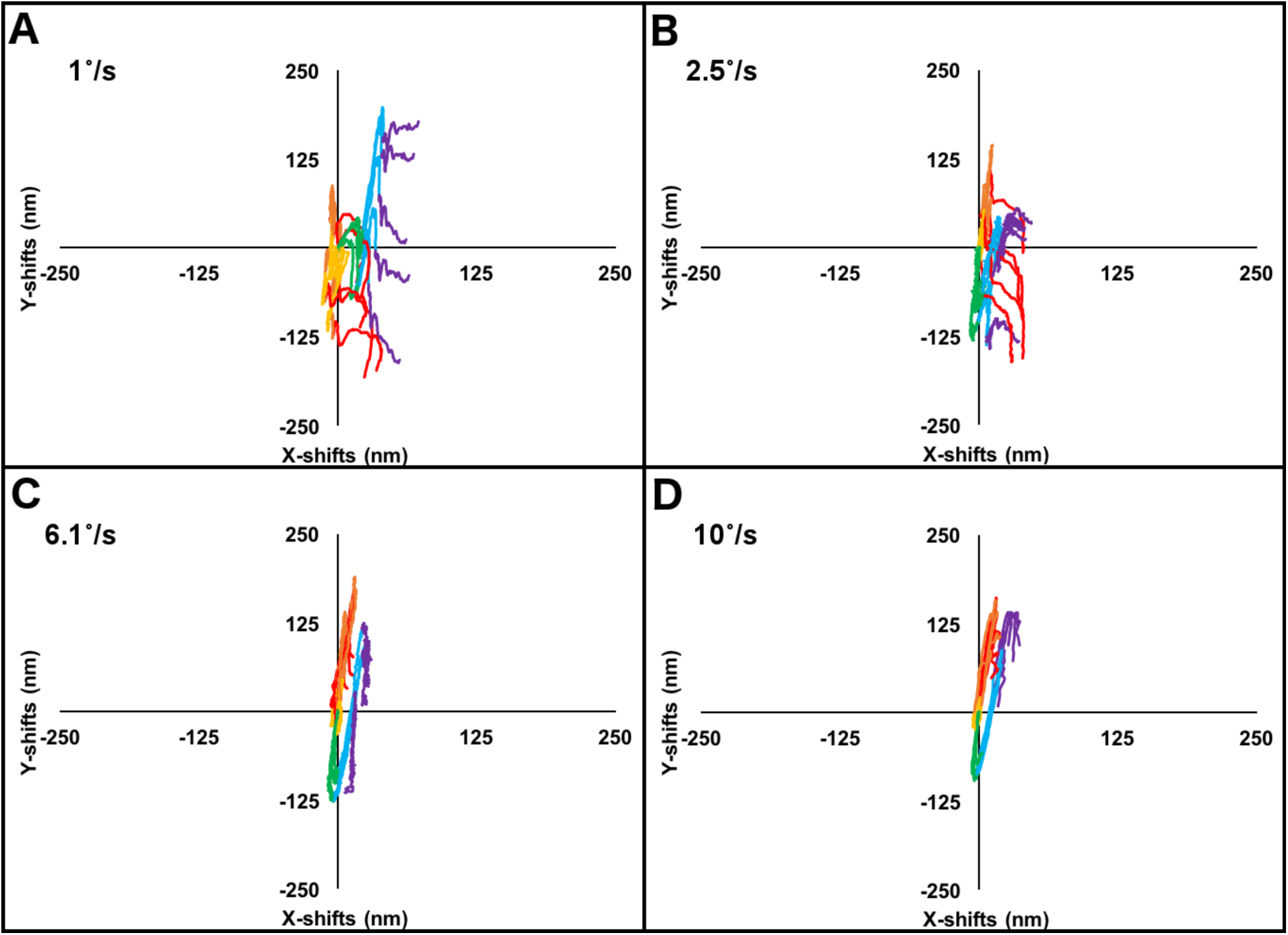
Stage eucentricity plots of XY shifts using the continuous tilting method at various tilting speeds (inset). The tilt axis is parallel to the x-axis. Each curve is depicted with a rainbow color gradient from red to violet, with red being −51° and violet being +51° tilt.

### Continuous-tilting method - quality of reconstructions

Tomograms obtained using the continuous-tilting method appeared by eye to be of similar quality as those obtained with conventional, slower methods (Figure 4). The two leaflets of the outer membrane lipid bilayer, which are typically 3-4 nm apart (Lewis and Engelman, 1983), were clearly visible (Figure 4A), suggesting a resolution of at least 4 nm. Other features, such as secretin pores (Figure 4B) (Chang et al., 2017), as well as side and top views of methyl-accepting chemotaxis protein (MCP) arrays (Figure 4C and 4D) (Briegel et al., 2009) also suggested ∼4 nm resolution. Unfortunately, the power spectra of individual images in the tilt-series exhibited no Thon rings, preventing CTF correction. We initially attributed the absence of information in the power spectrum to the low contrast of individual frames, but every attempt at motion-correction and averaging also failed to produce Thon rings (data not shown). To investigate why, we collected high-dose continuous tilt-series of cross gratings. Power spectra of the aligned images revealed partial Thon rings along the tilt axis, but rapid loss of information perpendicular to the tilt axis (Figure 5A). Varying the tilting speed produced the same effect. Averaging frames at different tilt angles and varying the number of frames included in averaged power spectra, down to 1° overall tilted angle, also produced power spectra with the same pattern. These observations suggest a vibration orthogonal to the tilt axis, most likely caused by the operation of the α tilt motor of the stage. This pattern suggests a vibration orthogonal to the tilt axis, most likely caused by the operation of the α tilt motor of the stage. Given these findings, the continuous-tilting method is only suitable for projects requiring low (∼4 nm) resolution.

**Figure 4.**
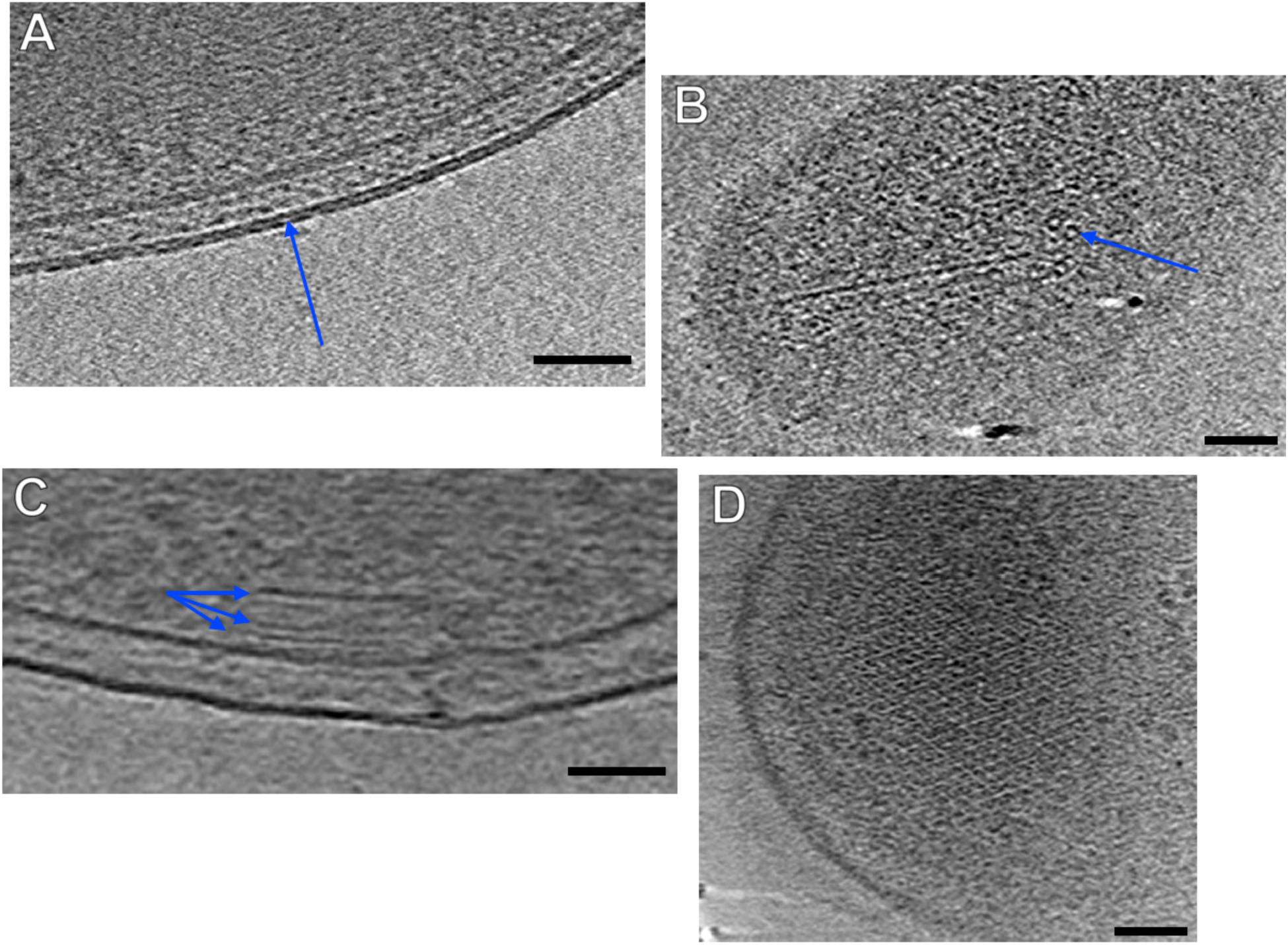
Observable features in several tomograms of *Bdellovibrio bacteriovorus* collected using the continuous tilting method. A) Double leaflets of the outer membrane lipid bilayer. B) Secretin pores, C) Side-view of MCP array. D) Top view of MCP array (Scale bars: 50nm).

**Figure 5.**
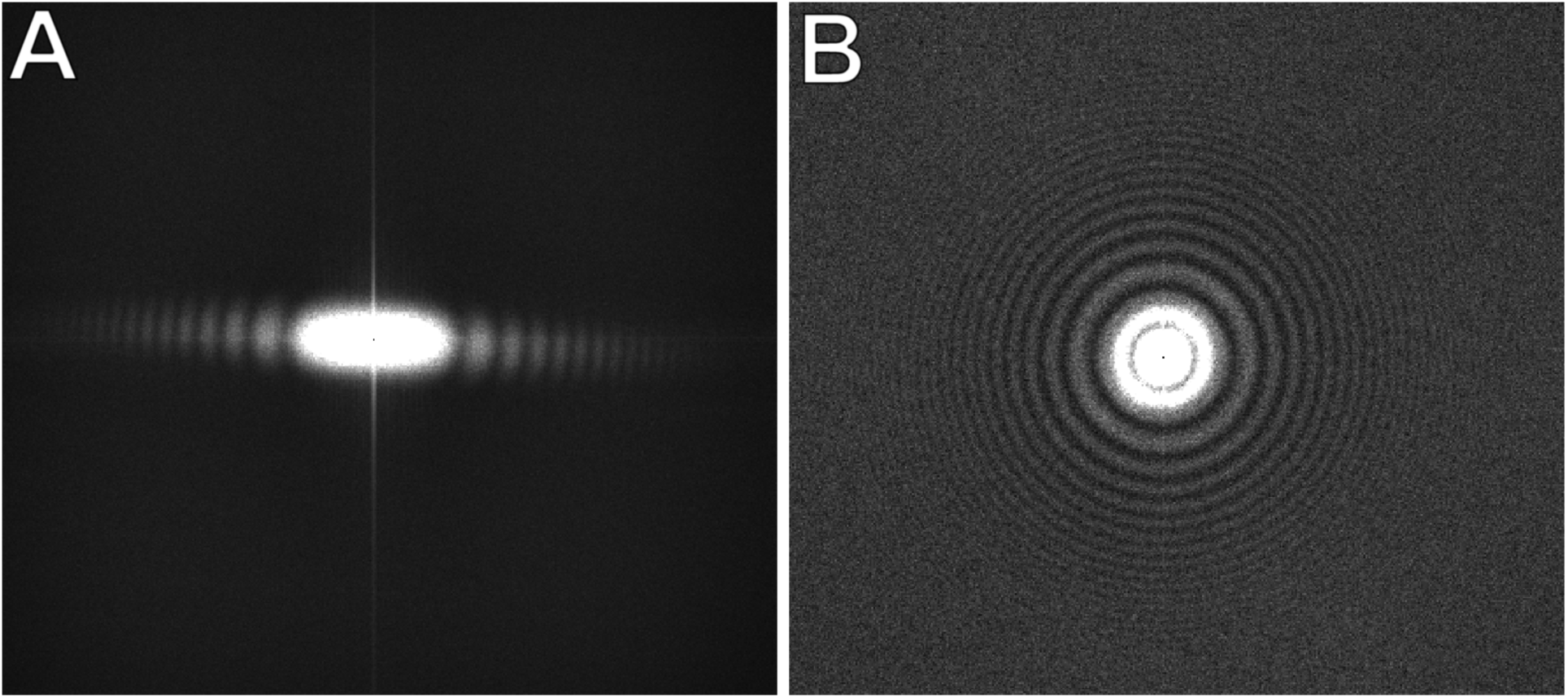
A) Power spectrum of high-dose continuous tilt-series taken of cross-gratings to show partial Thon rings along the tilt axis. B) Power spectrum of 0 degree tilt angle after fast-incremental tilting (tilting to 0 and immediately unblanking beam with no added delay time). Nyquist frequency is depicted at the edge of each image.

### Fast-incremental method - speed

We next developed a fast-incremental method, which solves the stage motor vibration problem by stopping the stage at discrete tilt angles. This method is depicted in the following scheme:

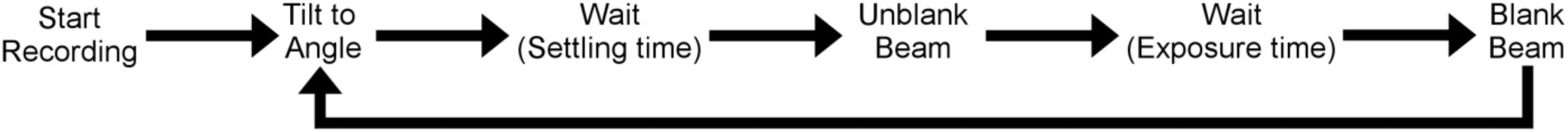

With a blanked electron beam, the camera is set to begin recording a long movie. The stage is then tilted to the first tilt angle in the tilt-scheme. After a specific settling time, defined as a delay time to allow the stage to settle after tilting, the beam is unblanked and the sample exposed for a preset exposure time. The beam is then blanked and the stage tilted to the next angle. This loop is repeated for each tilt angle in the tilt-scheme. We first characterized the settling time: how long does the stage need to settle after tilting to minimize vibration? We tested this by tilting the stage from −3° to 0°, varying the settling time from 0 ms to up to 5000 ms, and unblanking. After examining each resulting power spectrum for signs of vibration, we found that even with a settling time of 0 ms, the power spectrum displayed full Thon rings and no evidence of vibration (Figure 5B), suggesting that stage vibration we observed in continuous-tilting is associated with stage motion. We therefore proceeded with no settling time for all subsequent fast-incremental tilt-series collected. (Note that drift remains, but it can be compensated for by motion-correcting all the frames collected at each tilt angle.)

We then tested the speed of tilt-series acquisition using the fast-incremental method by recording tilt-series using three common tilt-series collection schemes: unidirectional, bi-directional and dose symmetric (Table 2). The total record time refers to the total time spent performing the tilt-schemes, while the total time per tilt-series refers to the total time required to re-gain microscope control. As expected, collection schemes which involved more tilting or a greater number of exposures resulted in a larger number of total frames and correspondingly longer times. We were surprised to find that the lock-out time for fast-incremental schemes is much shorter than those of the continuous-tilting method, even with tilt-series containing more frames. This is most likely due to the fact that most of the frames are blank, reducing the total amount of data needing to be stored.

**Table 2.** Time taken to collect fast-incremental tilt series. Files were saved without gain normalization in MRC format.

| Collection Scheme | Tilt Increment | Total number of frames | Exposure time per tilt | Total record time | Total time per ts |
| --- | --- | --- | --- | --- | --- |
| Unidirectional | 3° | 1050 | 1 sec | 97 sec | 2.6 min |
| Bidirectional | 3° | 950 | 1 sec | 89 sec | 2.7 min |
| Bidirectional | 1° | 3440 | 0.7 sec | 190 sec | 6 min |
| Dose Symmetric | 3° | 1450 | 1 sec | 138 sec | 4 min |

### Fast-incremental method - How eucentric is the stage?

We next characterized the eucentricity of the stage by plotting the XY translations required to align all the images of a tilt-series. Since we were able to see Thon rings in the power spectrum of individual images, we also measured the defocus for each image as an indicator of Z change throughout the tilt series. We characterized stage eucentricity for all 3 collection schemes: unidirectional, bidirectional, and dose symmetric (Figure 6A, 6C, and 6E). For all three collection schemes, most of the movement occurred perpendicular to the tilt axis and was remarkably consistent. Both unidirectional and bidirectional schemes exhibited larger shifts at tilt angles beyond +/- 40° (Figure 6A and 6C). Larger shifts were seen in Z, as the defocus changed by about 2-3 µm throughout the tilt series (Figure 6B, 6D, and 6F). This behavior could be partly due to a lateral offset between the optical axis and the tilt axis. The dose symmetric scheme exhibited the smallest overall shifts in both Y and Z (Figure 6E and 6F), which suggests that tilting back and forth may compensate for additive backlashes present in the other tilt-schemes. While overall the stage movements in the fast-incremental method were greater than those in the continuous-tilting method, the patterns were more predictable. This predictability should allow pre-calibration of the image movement prior to data collection (Ziese et al., 2002). The calibrated shifts could then be applied during data collection to compensate for sample movement in x, y, and z and greatly reduce field of view loss and defocus variation with the fast-incremental method.

**Figure 6.**
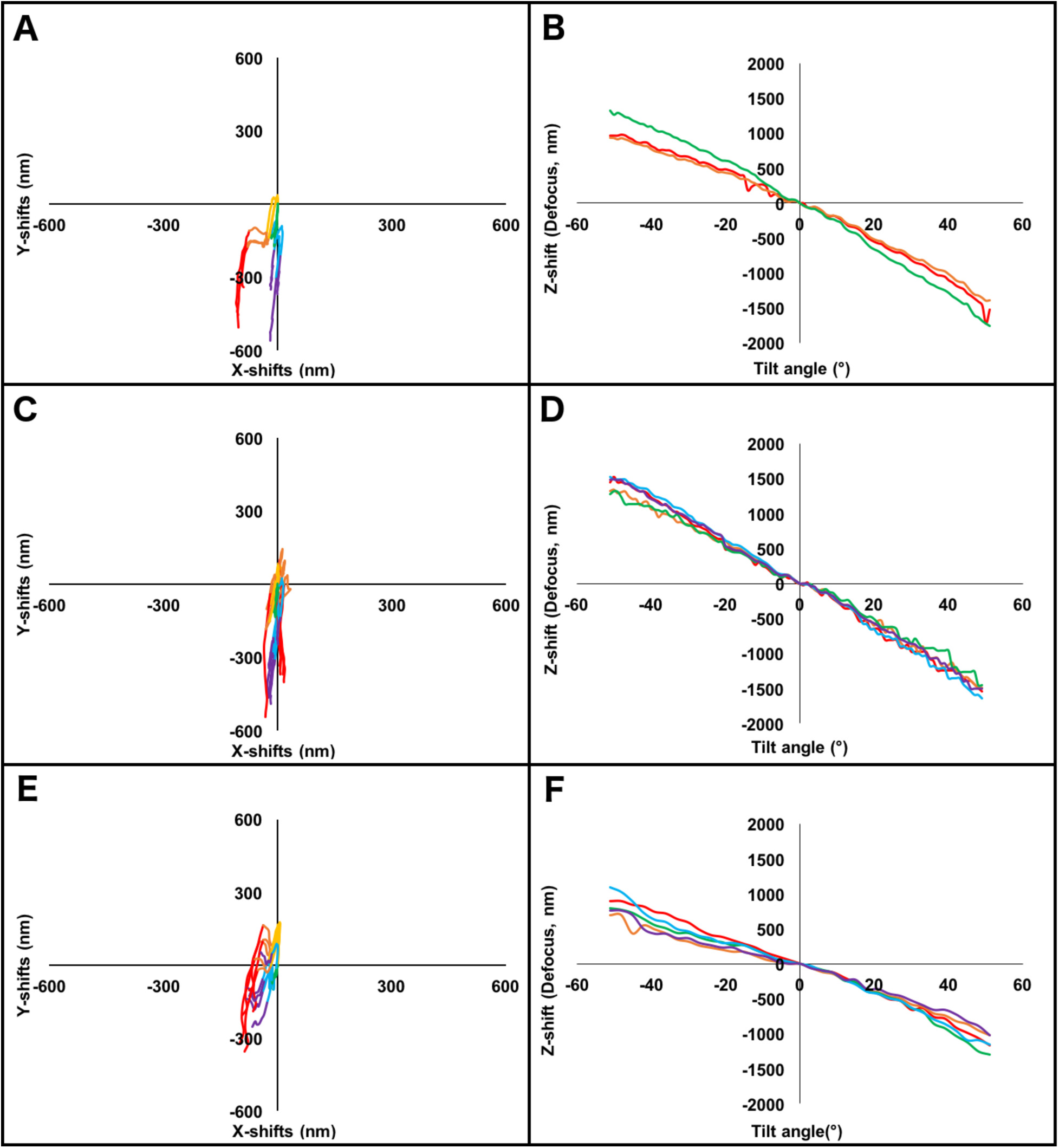
Stage eucentricity displayed as plots of X, Y, and Z shifts using the fast-incremental method and various collection schemes: A) Unidirectional XY shifts, −51° to +51°, 1° increment B) Z shifts for the data in A. C) Bidirectional XY shifts from −18° to ±52°, 1° increment. D) Z shifts for the data in C. E) Dose Symmetric XY shifts to ±51°, 3° increment. F) Z shifts for the data in E. The tilt axis is parallel to the x-axis. Each curve is depicted with a rainbow color gradient from red to violet, with red being −51° and violet being +51° tilt. Note that the scale here is 2.4x larger than depicted in Fig. 2.

### Fast-incremental tilting method - How does the quality of reconstructions compare to conventional ECT?

To assess the quality of the tomograms produced by the fast-incremental method, we collected tilt-series of *Bdellovibrio bacteriovorus*. We observed small (7-8 nm width) barrel-shaped proteins in the cytoplasm (Figure 7A) which could be GroEL, a 65 kDa chaperone found in many bacterial species (Kohda et al., 2000; Zeilstra-Ryalls et al., 1991). The double leaflets of both outer and inner membrane lipid bilayer were also clearly visible, as were side (Figure 7B) and top views (Figure 7C) of MCP arrays. A subtomogram average of 300 particles of an MCP array clearly revealed individual MCP dimers (Figure 7D), suggesting a resolution of at least 2.5 nm. These results are comparable to our previous work (Briegel et al., 2012), and suggest that the fast-incremental method is able to deliver similar resolutions as conventional ECT methods.

**Figure 7.**
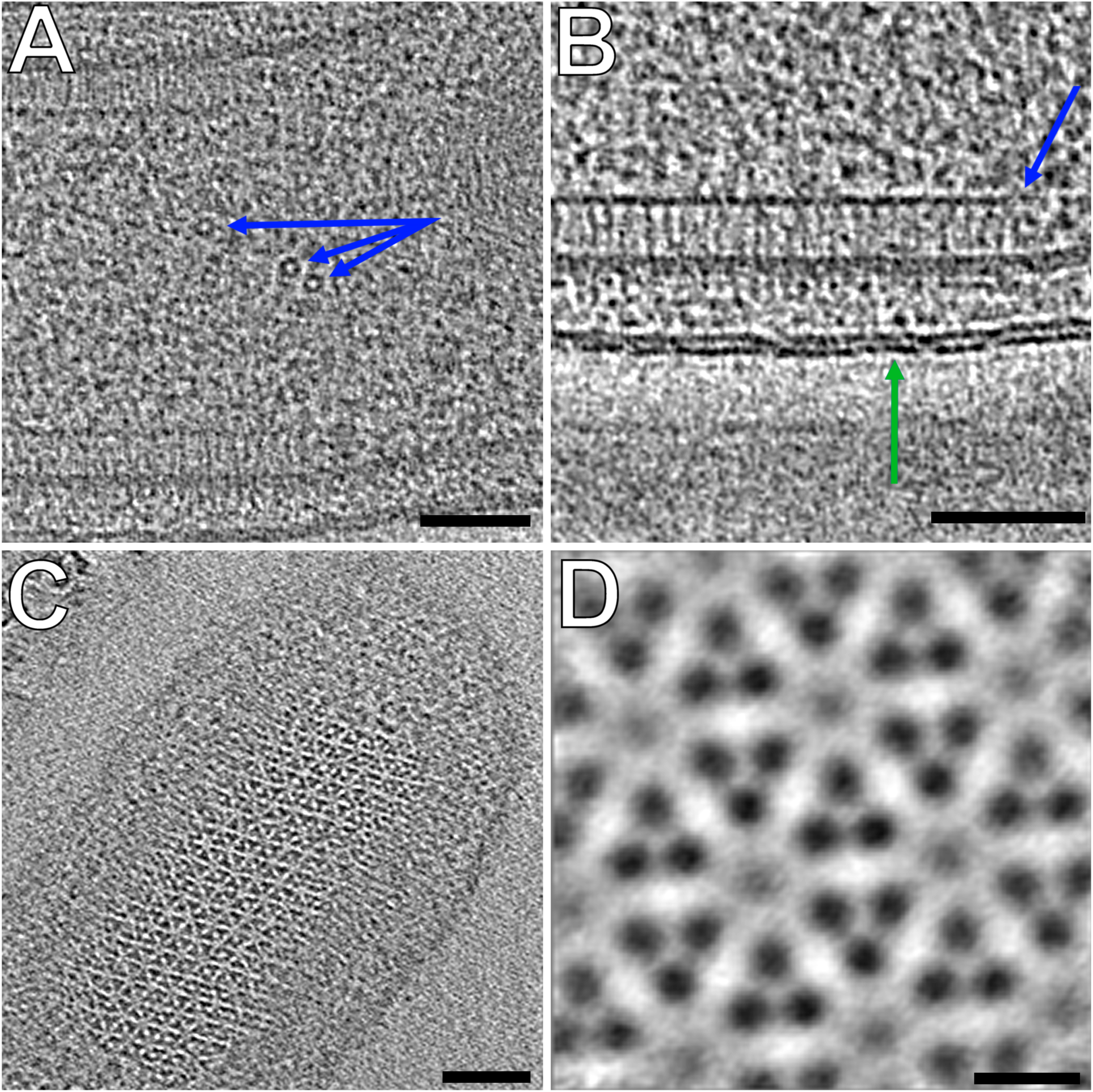
A, B and C) Observable features in a tomogram of *Bdellovibrio bacteriovorus* collected using the fast-incremental method, bi-directional scheme with 1° tilt increment. (Scale bars: 50nm). A) Side view of MCP array (blue arrow) and double leaflets of the outer membrane lipid bilayer (green arrow). B) Barrel-shaped protein complexes (blue arrows) C) Top view of MCP array D) Subtomogram average of MCP array shown in C (Scale bar: 5nm).

### Fast-incremental method - Potential to become the method of choice for ECT?

To assess whether the fast-incremental method is a good candidate to replace traditional ECT, we asked how much microscope time could be saved if it were applied to a conventional project that involved subtomogram averaging. To test this, we recorded 50 tilt-series of *B. bacteriovorus* using the fast-incremental method. Using a bidirectional tilt-scheme and 1° tilt increment, 7.5 mins per tilt-series were required: approximately 40 s were taken to find each target, 45 s to tune eucentric height, 5 s to autofocus and 6 min to perform the bidirectional collection scheme and regain control of the microscope (Table 2). Our conventional methods require similar times to find targets and tune the eucentric height, but much longer collection times due to tracking and to saving individual images, resulting in about 35 mins per tilt series, nearly 5 times longer than what we can currently achieve with the fast-incremental method.

While this improvement is significant, the fast-incremental method should soon become even faster. Most importantly, next-generation direct detectors will have much faster write speeds, eliminating latency. Target picking and eucentric height tuning could be automated. Finally, the tilt speed between exposures could be increased. Together, we estimate that the fast-incremental method could become 10-12 times faster than conventional ECT methods, allowing the acquisition of up to 500 tilt-series per day. If future stages can be built that do not vibrate when tilting, and next-generation cameras with much higher electron-counting speeds (Gatan’s K3 camera records 1500 frames per second) are used, the continuous-tilting method could allow tilt-series acquisition in just seconds. Methods to more reliably fine-tune eucentric height would result in a more accurately centered target, smaller field of view loss and less defocus variation. Once hundreds of tilt-series per day are collected, more reliable fiducial tracking software for tilt-series alignment will be needed. Finally, software that can automatically identify and pick objects for subtomogram averaging will allow the entire ECT and subtomogram averaging workflow to become automated.

The development of methods to collect tilt-series in just tens of seconds will obviously accelerate what we think of today as tomography projects, making entirely new kinds of structural studies possible, but it may also profoundly alter other cryo-EM workflows as well. Because for a given dose, a tilt-series provides more information than a single projection (the dose fractionation theorem, Hegerl and Hoppe, 1976) we expect future single particle workflows, for instance, to begin with a medium-dose (∼30 e^−^/Å^2^) projection image and end with a fast tilt-series. While only the initial, early-dose projections will likely be used in the final reconstruction, the tomograms could be used to identify potentially damaged particles at the air/water interface, improve per-particle defocus estimates, improve initial models, improve particle alignment, and disambiguate conformational changes from different orientations.

## Methods

### Bdellovibrio bacteriovorus cell growth and preparation for ECT

Host-dependent *B. bacteriovorus* cells were grown as previously described (Lambert and Sockett, 2008). First, wild type *Escherichia coli* cells were inoculated into 2 mL of LB media and incubated for 8 h at 37 °C with agitation. 1% YPSC agar was poured onto a plate and allowed to solidify as a bottom layer. 100 µl of *E. coli* starter culture was then mixed with a small volume of warm 0.6 % YPSC agar, poured onto the plate and allowed to solidify as a top layer. A small amount of previously prepared *B. bacteriovorus* glycerol stocks were pipetted at the very center of the plate and incubated upright at 29 °C until a lawn of E. coli growth appeared. The plate was then flipped upside-down and incubated for 4 additional days until a clear halo appeared at the center, indicating prey cells had been killed. The clear halo portion of agar was then transferred into 2 mL HEPES buffer supplemented with 10 mM CaCl_2_ and 150 µL of fresh *E. coli* culture, and incubated at 29 °C with agitation for 3 days. Suitable *B. bacteriovorus* cell growth and concentration were confirmed by negative stain and imaging on a Tecnai T12 electron microscope (FEI Company). 3 µl of cells mixed with 10nm gold beads were pipetted onto freshly glow-discharged Quantifoil copper R2/2 200 EM grid (Quantifoil Micro Tools GmbH) and plunge-frozen into a liquid ethane-propane mixture using an Vitrobot Mark-IV (Thermo Fisher Scientific).

### Electron tomography data collection

All tomographic tilt-series were collected in electron-counting mode using *SerialEM* software versions 3-7-0beta4 to 3-7-0beta10 (Mastronarde, 2003) on a Titan Krios (Thermo Fisher Scientific) equipped with a Gatan energy filter and a K2 Summit direct electron detector (Gatan). *B. bacteriovorus* tilt-series were collected using the continuous-tilting or fast-incremental methods with a total dosage of 100 e^−^/Å^2^ and defocus values ranging from −2 µm to −5 µm.

#### The continuous-tilting method

With the continuous-tilting method, the stage is continuously tilted in one direction as the camera simultaneously records and saves movie frames at a frame-rate of 20-40 frames per second (fps) using *SerialEM*’s TiltDuringRecord script command. This results in tilt-series containing hundreds to thousands of movie frames with very small angular intervals between each image, ranging from 0.1°/image to 0.25°/image, depending on tilting speed and camera framerate. For bi-directional tilt-schemes, two separate movies were collected for each direction and combined using IMOD’s *Newstack* command (Kremer et al., 1996). Stage tilting speeds were adjusted to meet the electron counting rate of the K2 Summit detector while maximizing SNR and minimizing radiation damage. Smaller pixel sizes deliver greater doses per A^2^, allowing for greater tilting speeds.

#### The fast-incremental method

We tested three types of tilt-schemes at 1°, 2°, and 3° tilt increments: the unidirectional scheme involves tilting the stage in a single direction, from −60° to +60°. The bidirectional scheme involves tilting the stage from −18° to +51° or +52°, then from −20° to −51° or −52°. Finally, the dose symmetric scheme was adapted from Hagen et al. to maximize high-resolution information (Hagen et al., 2017) by starting at a low tilt angle and tilting the stage back and forth, starting at 0 °, then 3°, −3°, 6°, −6°, and so on until −51° and +51°. All eucentricity measurements were done with 3-5 iterations for each tilt-scheme at different locations on the grid.

#### SerialEM scripts

During the fast-incremental method, camera recording in *SerialEM* disables the user interface and prevents microscope control. We therefore used a second *SerialEM* program to control the stage and beam. The latest version of SerialEM has removed this limitation.

### Electron tomography data processing

All tilt-series were processed and aligned using IMOD (Kremer et al., 1996). Defocus measurements were done using IMOD’s *Ctfplotter*, EMAN2 (Tang et al., 2007, p. 2) or *CTFFIND4* (Rohou and Grigorieff, 2015), and contrast-transfer function (CTF) correction done by phase inversion using IMOD’s *Ctfphaseflip*. Tomographic reconstructions by weighted back-projection were done using IMOD’s *Tilt*, while SIRT reconstructions were produced using Tomo3D (Agulleiro and Fernandez, 2011). Subtomogram averaging was performed using Dynamo software (Castaño-Díez et al., 2012).

#### Alignment of low contrast frames obtained with continuous-tilting

Because frames of continuous tilt-series have very low SNR, we developed *Neighbor-enhance*, a new script that increases SNR by averaging neighboring frames together. For each image in a raw tilt-series obtained with the continuous-tilting method, *Neighbor-enhance* uses IMOD functions to extract neighboring sets of frames. With the middle frame as a reference, each neighboring image is stretched by an amount equal to the ratio of the cosines of its tilt angle and of the tilt angle of the reference frame. Stretched, motion-corrected sets of frames are finally averaged. This averaging greatly improves SNR and facilitates fiducial picking so a fine transformation matrix can be calculated and applied to the raw low contrast tilt-series (Figure 2).

#### Automated removal of blank frames obtained with the fast-incremental method

The *removeblankframes* script first gain-normalizes, corrects defects, and runs IMOD’s *Ccderaser* to remove deviant pixels on a raw fast-incremental tilt-series. It then extracts electron counts from each image of the tilt-series and excludes those with low electron counts, indicating a blank frame. Finally, the remaining non-blank movie frames are extracted and motion-corrected using IMOD’s *Alignframes* to generate a final tilt stack for regular processing using a traditional workflow, such as IMOD’s *Etomo*.

*Both Neighbor-enhance* and *Removeblankframes* scripts are available for download at https://jensenlab.caltech.edu/.

## Data Deposition

Tilt-series and tomographic reconstructions were deposited in the Electron Microscopy Data Bank (EMDB accession codes: EMD-9260 and EMD-9261), and the Electron Microscopy Public Image Archive (EMPIAR accession codes: EMPIAR-10225 and EMPIAR-10226)

## Acknowledgements

We thank David N. Mastronarde for helpful suggestions and for providing comments on this manuscript. This work was supported by NIH grant GM122588 (to G.J.J.). Electron cryomicroscopy was done in the Beckman Institute Resource Center for Transmission Electron Microscopy at Caltech.

## References

Agulleiro, J.I., Fernandez, J.J., 2011. Fast tomographic reconstruction on multicore computers. Bioinformatics 27, 582–583. https://doi.org/10.1093/bioinformatics/btq692

Briegel, A., Li, X., Bilwes, A.M., Hughes, K.T., Jensen, G.J., Crane, B.R., 2012. Bacterial chemoreceptor arrays are hexagonally packed trimers of receptor dimers networked by rings of kinase and coupling proteins. PNAS. https://doi.org/10.1073/pnas.1115719109

Briegel, A., Ortega, D.R., Tocheva, E.I., Wuichet, K., Li, Z., Chen, S., Müller, A., Iancu, C.V., Murphy, G.E., Dobro, M.J., Zhulin, I.B., Jensen, G.J., 2009. Universal architecture of bacterial chemoreceptor arrays. PNAS pnas.0905181106. https://doi.org/10.1073/pnas.0905181106

Briggs, J.A., 2013. Structural biology in situ—the potential of subtomogram averaging. Current Opinion in Structural Biology, Theory and simulation / Macromolecular assemblies 23, 261–267. https://doi.org/10.1016/j.sbi.2013.02.003

Castaño-Díez, D., Kudryashev, M., Arheit, M., Stahlberg, H., 2012. Dynamo: a flexible, user-friendly development tool for subtomogram averaging of cryo-EM data in high-performance computing environments. Journal of structural biology 178, 139–151.

Chang, Y.-W., Kjær, A., Ortega, D.R., Kovacikova, G., Sutherland, J.A., Rettberg, L.A., Taylor, R.K., Jensen, G.J., 2017. Architecture of the *Vibrio cholerae* toxin-coregulated pilus machine revealed by electron cryotomography. Nature Microbiology 2, 16269. https://doi.org/10.1038/nmicrobiol.2016.269

Chang, Y.-W., Rettberg, L.A., Treuner-Lange, A., Iwasa, J., Søgaard-Andersen, L., Jensen, G.J., 2016. Architecture of the type IVa pilus machine. Science 351, aad2001.

Hagen, W.J.H., Wan, W., Briggs, J.A.G., 2017. Implementation of a cryo-electron tomography tilt-scheme optimized for high resolution subtomogram averaging. Journal of Structural Biology, SI:Electron Tomography 197, 191–198. https://doi.org/10.1016/j.jsb.2016.06.007

Hegerl, R., Hoppe, W., 1976. Influence of Electron Noise on Three-dimensional Image Reconstruction. Zeitschrift für Naturforschung A 31, 1717–1721. https://doi.org/10.1515/zna-1976-1241

Hu, B., Morado, D.R., Margolin, W., Rohde, J.R., Arizmendi, O., Picking, W.L., Picking, W.D., Liu, J., 2015. Visualization of the type III secretion sorting platform of Shigella flexneri. PNAS 112, 1047–1052. https://doi.org/10.1073/pnas.1411610112

Kohda, J., Kondo, A., Teshima, T., Fukuda, H., 2000. Development of Efficient Protein Refolding Systems Using Chaperonins, in: Endo, I., Nagamune, T., Katoh, S., Yonemoto, T. (Eds.), Progress in Biotechnology, Bioseparation Engineering. Elsevier, pp. 119–124. https://doi.org/10.1016/S0921-0423(00)80023-4

Kremer, J.R., Mastronarde, D.N., McIntosh, J.R., 1996. Computer Visualization of Three-Dimensional Image Data Using IMOD. Journal of Structural Biology 116, 71–76. https://doi.org/10.1006/jsbi.1996.0013

Lambert, C., Sockett, R.E., 2008. Laboratory maintenance of Bdellovibrio. Current protocols in microbiology 9, 7B. 2.1–7B. 2.13.

Lewis, B.A., Engelman, D.M., 1983. Lipid bilayer thickness varies linearly with acyl chain length in fluid phosphatidylcholine vesicles. Journal of Molecular Biology 166, 211–217. https://doi.org/10.1016/S0022-2836(83)80007-2

Mastronarde, D.N., 2005. Automated electron microscope tomography using robust prediction of specimen movements. Journal of Structural Biology 152, 36–51. https://doi.org/10.1016/j.jsb.2005.07.007

Mastronarde, D.N., 2003. SerialEM: A Program for Automated Tilt Series Acquisition on Tecnai Microscopes Using Prediction of Specimen Position. Microscopy and Microanalysis 9, 1182–1183. https://doi.org/10.1017/S1431927603445911

Murphy, G.E., Leadbetter, J.R., Jensen, G.J., 2006. In situ structure of the complete Treponema primitia flagellar motor. Nature 442, 1062.

Nickell, S., Förster, F., Linaroudis, A., Net, W.D., Beck, F., Hegerl, R., Baumeister, W., Plitzko, J.M., 2005. TOM software toolbox: acquisition and analysis for electron tomography. Journal of Structural Biology 149, 227–234. https://doi.org/10.1016/j.jsb.2004.10.006

Rohou, A., Grigorieff, N., 2015. CTFFIND4: Fast and accurate defocus estimation from electron micrographs. J. Struct. Biol. 192, 216–221. https://doi.org/10.1016/j.jsb.2015.08.008

Suloway, C., Shi, J., Cheng, A., Pulokas, J., Carragher, B., Potter, C.S., Zheng, S.Q., Agard, D.A., Jensen, G.J., 2009. Fully automated, sequential tilt-series acquisition with Leginon. Journal of structural biology 167, 11–18.

Tang, G., Peng, L., Baldwin, P.R., Mann, D.S., Jiang, W., Rees, I., Ludtke, S.J., 2007. EMAN2: An extensible image processing suite for electron microscopy. Journal of Structural Biology, Software tools for macromolecular microscopy 157, 38–46. https://doi.org/10.1016/j.jsb.2006.05.009

Zeilstra-Ryalls, J., Fayet, O., Georgopoulos, C., 1991. The Universally Conserved GroE (Hsp60) Chaperonins. Annual Review of Microbiology 45, 301–325. https://doi.org/10.1146/annurev.mi.45.100191.001505

Zheng, S.Q., Sedat, J.W., Agard, D.A., 2010. Chapter Twelve - Automated Data Collection for Electron Microscopic Tomography, in: Jensen, G.J. (Ed.), Methods in Enzymology, Cryo-EM Part A Sample Preparation and Data Collection. Academic Press, pp. 283–315. https://doi.org/10.1016/S0076-6879(10)81012-2

Ziese, U., Janssen, A.H., Murk, J.-L., Geerts, W.J.C., Van der Krift, T., Verkleij, A.J., Koster, A.J., 2002. Automated high-throughput electron tomography by pre-calibration of image shifts. Journal of Microscopy 205, 187–200.

